# Relationship between impulsivity, uncontrolled eating and body mass index: a hierarchical model

**DOI:** 10.1101/348821

**Authors:** Isabel Garcia-Garcia, Selin Neseliler, Filip Morys, Mahsa Dadar, Yvonne H.C. Yau, Stephanie G. Scala, Yashar Zeighami, Natalie Sun, D. Louis Collins, Uku Vainik, Alain Dagher

**Affiliations:** Montreal Neurological Institute, McGill University, Montréal, Canada; Institute of Psychology, University of Tartu, Estonia

**Keywords:** Obesity, MRI, impulsivity, weight gain, Uncontrolled Eating

## Abstract

**Background:** Impulsivity increases the risk for obesity and weight gain. However, the precise role of impulsivity in the aetiology of overeating behavior and obesity is currently unknown. Here we examined the relationships between personality-related measures of impulsivity, Uncontrolled Eating, BMI, and longitudinal weight changes. Additionally, we analyzed the associations between general impulsivity domains and brain cortical thickness to elucidate brain vulnerability factors related to weight gain.

**Methods:** Students in their first year of university - a risky period for weight gain - completed questionnaire measures of impulsivity and eating behavior at the beginning (N = 2318) of the school year. We also collected their weight at the end of the term (N = 1197). Impulsivity was divided into factors stress reactivity, reward sensitivity and lack of self-control. Using structural equation models, we tested the plausibility of a hierarchical relationship, in which impulsivity traits were associated with Uncontrolled Eating, which in turn predicted BMI and weight change. 71 participants underwent T1-weighted MRI to investigate the correlation between impulsivity and cortical thickness.

**Results:** Impulsivity traits showed positive correlations with Uncontrolled Eating. Higher scores in Uncontrolled Eating were in turn associated with higher BMI. None of the impulsivity-related measurements nor Uncontrolled Eating were correlated with longitudinal weight gain. Higher stress sensitivity was associated with increased cortical thickness in the superior temporal gyrus. Lack of self-control was positively associated with increased thickness in the superior medial frontal gyrus. Finally, higher reward sensitivity was associated with lower thickness in the inferior frontal gyrus.

**Conclusion:** The present study provides a comprehensive characterization of the relationships between different facets of impulsivity and obesity. We show that differences in impulsivity domains might be associated with BMI via Uncontrolled Eating. Our results might inform future clinical strategies aimed at fostering self-control abilities to prevent and/or treat unhealthy weight gain.

## Introduction

The increase in the incidence of obesity in the past 50 years can be attributed to overeating in response to abundant and inexpensive calories (1). Obesity is also heritable, and most of the implicated genes appear to be expressed in the central nervous system (2). As such, obesity can be thought of as resulting from an interaction between a brain-based endophenotype and a disease-promoting environment (3). Endophenotypes are intermediate phenotypes that link latent biological processes to observable outcomes (4,5). The neurobehavioral endophenotypes associated with obesity can be broadly categorized as domain-general and eating-specific (6).

One domain-general endophenotype is impulsivity (7), the tendency to act without full consideration of the consequences (8). It can be subdivided into three domains that align with personality factors: 1) low *conscientiousness*, reflecting self-control, or, in other terms, premeditation and perseverance; 2) *neuroticism*, a reflection of an individual’s sensitivity to stress and aversive events, also referred to as negative urgency, or the tendency to act impulsively when distressed, and 3) *extraversion*, reflecting sensitivity to rewards and sensation seeking (8–10). Enhanced impulsivity is characteristic of different neuropsychiatric conditions, such as pathological gambling, substance abuse and attention deficit hyperactivity disorder (11,12). Additionally, it might constitute a risk factor for obesity and overeating patterns (13–15).

Meta-analytic studies have shown that impulsivity shows weak positive correlations with BMI across independent studies (16). The effects of impulsivity on BMI, however, seem to be largely heterogeneous (16). One of the reasons for this heterogeneity could be explained by different impulsivity measures and domains used across studies (6). Another possibility is that the relation between personality measures of impulsivity and BMI might be mediated by individually-varying eating-specific impulsivity characteristics (17). In terms of brain correlates of impulsivity, it seems that they overlap with brain correlates of high BMI, which suggests that there is a relationship between obesity and impulsivity that might stem from the brain. For example, both high impulsivity and BMI are positively related to the volume of the striatum (18,19), or negatively related to the volume of the prefrontal and orbitofrontal cortex (20,21).

Eating-specific constructs associated with a high BMI include emotional eating (22), disinhibited eating (23), and power of food (24). Scores on all these questionnaires consistently and strongly correlate with BMI (25) and with each other (26). For this reason, in previous research we have proposed that different eating behavior questionnaires depict a common underlying latent factor, labelled uncontrolled eating (25,27). Preliminary results show that uncontrolled eating is associated with brain alterations from functional MRI studies, specifically higher brain activity in response to food cues or at rest in the cerebellum, and lower brain activity in the prefrontal cortex (28). While both impulsivity traits and uncontrolled eating have been linked to increased BMI (17), the relationships between different impulsivity domains, uncontrolled eating and obesity remain to be tested. A hypothesis here is that some impulsivity traits might be associated with eating behaviors, which in turn will be linked to BMI (17). However, comprehensive models have not been built for different impulsivity domains and eating questionnaires and have not been applied to the prediction of weight changes yet.

So far, most analyses have been conducted on cross-sectional datasets making it difficult to say if impulsivity is a risk factor or a consequence of obesity (28). The freshman year of university is an ideal time period to test these hypotheses both cross-sectionally and longitudinally. During this time, students transition into a new environment with access to similar food and exercise options, allowing underlying vulnerability to express itself. Weight gain often happens in this short period of time and affects approximately 50-60% of students (29,30). Eating specific behaviors that revolve around the concept of impulsivity have been most commonly studied as risk factors for weight gain. The results, however, have been contradictory (31–33).

In the brain, a high BMI has been associated with lower gray matter volume and cortical thickness in areas such as the medial prefrontal cortex extending to the orbitofrontal cortex, and temporal pole (21,34,35). However, it is possible that other brain structures involved in processes like impulsivity might be indirectly associated with weight gain. Identifying those regions might pave the way for new studies to investigate links between brain function and maladaptive eating patterns.

In the current study, we present a comprehensive examination of the relationships between impulsivity, uncontrolled eating, BMI and weight changes. We tested the hypothesis that the relations between general impulsivity, uncontrolled eating, BMI and longitudinal weight changes might be plausibly represented in a hierarchical structural equation model (SEM). In a sub-sample of participants, we additionally examined the relationship between domain-general impulsivity variables and brain structure. This post-hoc analysis was performed in order to extend our behavioral findings and identify new brain structures that might be relevant to weight status in an indirect manner.

## Methods and Materials

### Participants

Participants were first year McGill university students, at least 18 years of age, recruited via an advertisement sent to the incoming class electronic mailing list. Participants provided their consent online and data were collected using the online survey tool LimeSurvey (https://www.limesurvey.org) over three consecutive years. The study was approved by the Montreal Neurological Institute Research Ethics Board. Participants for the brain imaging experiment gave additional written consent before participating.

### Questionnaires

Participants filled out the following online questionnaires in the fall semester: “Big Five” personality dimensions (Openness/ Imagination, Conscientiousness, Agreeableness, Extraversion, and Neuroticism) from the International Personality Item Pool (IPIP) (36), two subscales (lack of perseverance and sensation seeking) of the UPPS Impulsive Behavior Scale (37), Cohen’s Perceived Stress Scale (PSS) (38), and Rosenberg Self Esteem Scale (39), as well as eating specific questionnaires such as the disinhibition subscale of Three-Factor Eating Questionnaire (TFEQ) (23), emotional eating subscale of Dutch Eating Behavior Questionnaires (DEBQ) (22), and all subscales of Power of Food Scale (PFS) (24) (Table 1). Participants also reported their height and weight at two timepoints – together with online questionnaires during the initial assessment in the fall semester, and during a follow-up in the spring semester. Only a subset of participants reported their height and weight during the follow-up (N=1145). Our questionnaires included two “catch” questions and three “catch-match” questions as a measure of the level of participants’ attention in completing the survey. Participants with total catch scores above three were excluded, resulting in a sample of 2318 participants (Males N= 750; Females N= 1538). After completion of the questionnaires during the initial assessment and after the follow-up, a subset of participants were asked to visit our laboratory to have their BMI measured. Height and weight were measured from 333 participants in the fall, and from 209 participants in the spring semester (N =115 overlap with the fall group) using a medical scale and a stadiometer. Self-reported BMI was highly correlated with measured BMI in the fall (r = 0.91, p<10^−5^) and in the spring (r = 0.92, p<10^−5^). We replaced the reported BMI with measured BMI in the analyses for higher accuracy. BMI was further residualized for age, sex and a covariate to account for whether it was derived from self-report or measured. Brain imaging was conducted on a subset of participants selected from this sample in the fall (N=71 participants, N=69 of them were included in the cortical thickness analysis, see below).

**Table 1.**
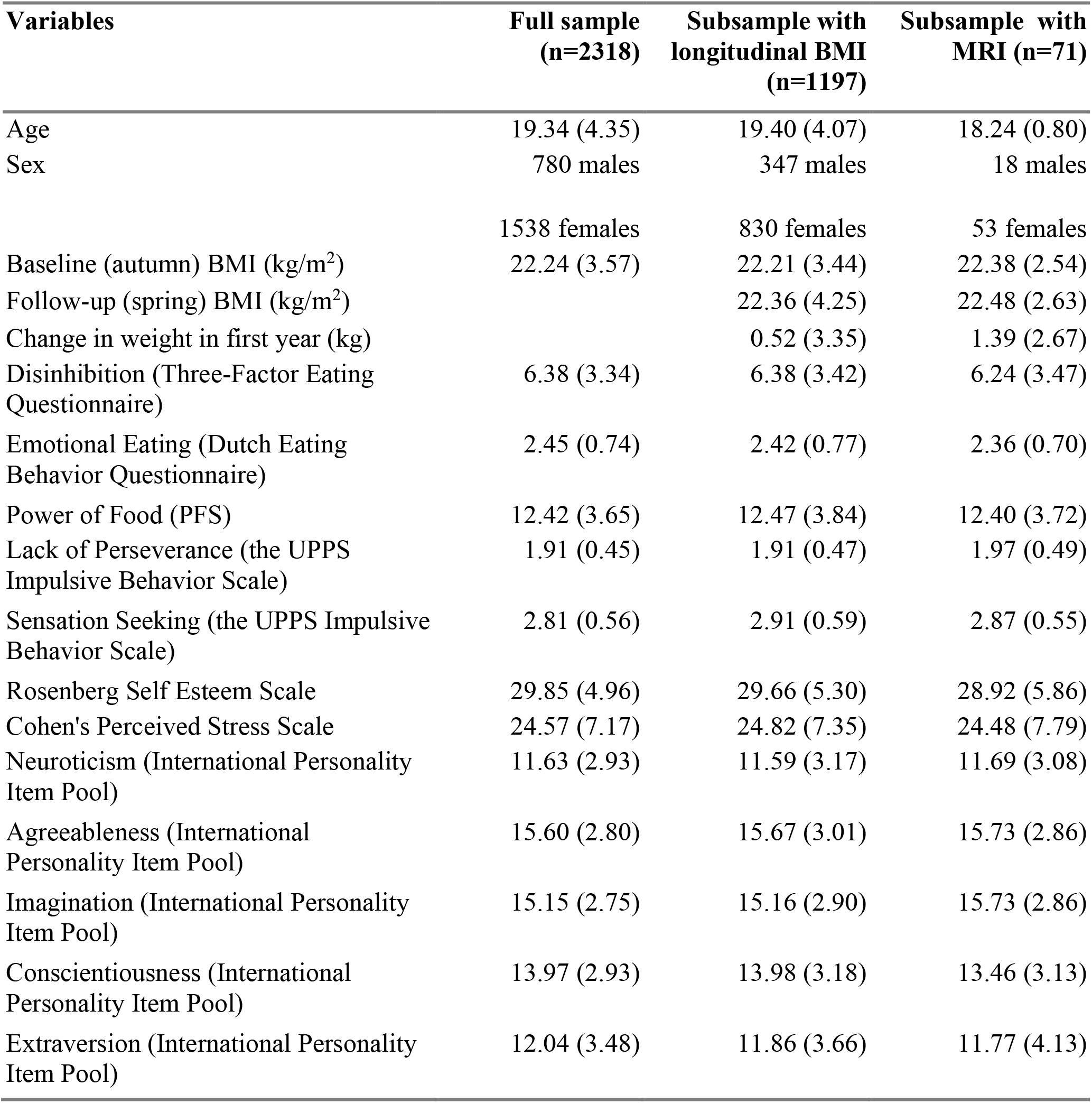
Descriptive statistics of the study variables. Numbers in parentheses indicate standard deviation.

| Variables | Full sample<br>(n=2318) | Subsample with<br>longitudinal BMI<br>(n=1197) | Subsample with<br>MRI (n=71) |
| --- | --- | --- | --- |
| Age | 19.34 (4.35) | 19.40 (4.07) | 18.24 (0.80) |
| Sex | 780 males<br>1538 females | 347 males<br>830 females | 18 males<br>53 females |
| Baseline (autumn) BMI (kg/m<sup>2</sup>) | 22.24 (3.57) | 22.21 (3.44) | 22.38 (2.54) |
| Follow-up (spring) BMI (kg/m<sup>2</sup>) |  | 22.36 (4.25) | 22.48 (2.63) |
| Change in weight in first year (kg) |  | 0.52 (3.35) | 1.39 (2.67) |
| Disinhibition (Three-Factor Eating Questionnaire) | 6.38 (3.34) | 6.38 (3.42) | 6.24 (3.47) |
| Emotional Eating (Dutch Eating Behavior Questionnaire) | 2.45 (0.74) | 2.42 (0.77) | 2.36 (0.70) |
| Power of Food (PFS) | 12.42 (3.65) | 12.47 (3.84) | 12.40 (3.72) |
| Lack of Perseverance (the UPPS Impulsive Behavior Scale) | 1.91 (0.45) | 1.91 (0.47) | 1.97 (0.49) |
| Sensation Seeking (the UPPS Impulsive Behavior Scale) | 2.81 (0.56) | 2.91 (0.59) | 2.87 (0.55) |
| Rosenberg Self Esteem Scale | 29.85 (4.96) | 29.66 (5.30) | 28.92 (5.86) |
| Cohen's Perceived Stress Scale | 24.57 (7.17) | 24.82 (7.35) | 24.48 (7.79) |
| Neuroticism (International Personality Item Pool) | 11.63 (2.93) | 11.59 (3.17) | 11.69 (3.08) |
| Agreeableness (International Personality Item Pool) | 15.60 (2.80) | 15.67 (3.01) | 15.73 (2.86) |
| Imagination (International Personality Item Pool) | 15.15 (2.75) | 15.16 (2.90) | 15.73 (2.86) |
| Conscientiousness (International Personality Item Pool) | 13.97 (2.93) | 13.98 (3.18) | 13.46 (3.13) |
| Extraversion (International Personality Item Pool) | 12.04 (3.48) | 11.86 (3.66) | 11.77 (4.13) |

### Structural Equation Models

We performed a series of SEMs to analyze the relationships between Uncontrolled Eating, the three impulsivity traits defined above, and weight gain. First, we tested the hypothesis that Uncontrolled Eating, stress reactivity, lack of self-control and reward sensitivity can be considered separable latent constructs. We built a four-dimension model (Model 1, Figure 1A), with Uncontrolled Eating, stress reactivity, lack of self-control and reward sensitivity modelled as latent constructs. Uncontrolled Eating (UE) was defined by the scores in Disinhibition, Power of Food and Emotional Eating, following previous work from our group (27,28). Stress reactivity was the latent variable that resulted from Perceived Stress Scale (PSS), Rosenberg self-esteem questionnaire and Neuroticism (IPIP). Lack of Self Control was a latent factor emerging from the observable variables Lack of Perseverance (UPPS) and Conscientiousness (IPIP). Finally, reward Sensitivity was formed by Extraversion (IPIP) and Sensation Seeking (UPPS). This model was compared to a simpler one-dimension model (Model 2, Figure 1B), where all the observable factors were fit into a single latent variable.

**Figure 1.**
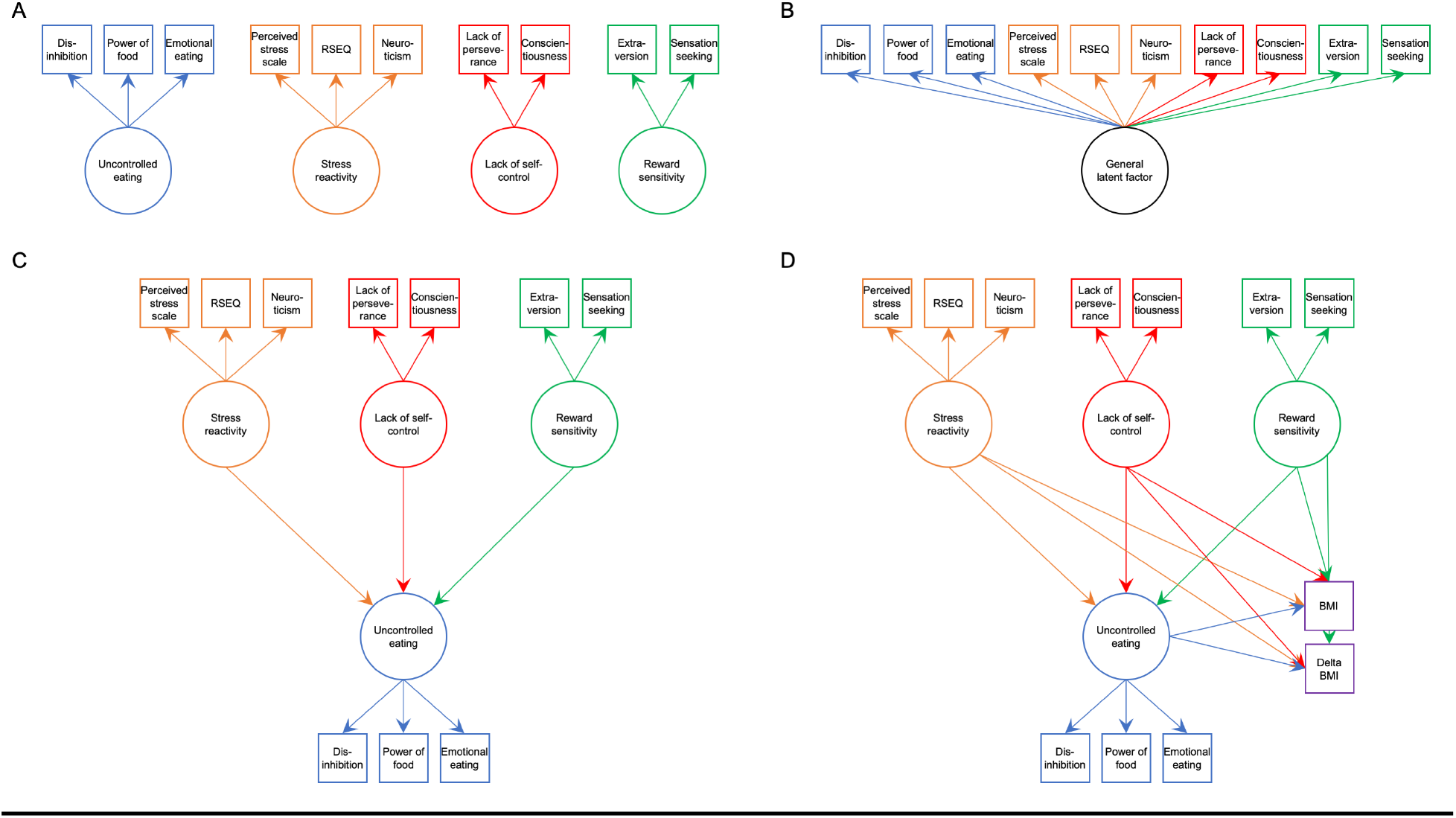
Structural equation models used in the study. **A** Model 1 representing 4 distinct latent variables; **B** Model 2 pooling all measures into 1 latent variable; **C** Model 3 representing a hierarchical model where stress reactivity, lack of self-control and reward sensitivity all influence Uncontrolled Eating; **D** Model 4 where all 4 latent variables influence BMI and BMI change. RSEQ – Rosenberg self-esteem questionnaire. BMI – body mass index.

We also built a hierarchical version of the four-dimension model (Model 3, Figure 1C), where we examined the plausibility of a layered model structure, inspired by the watershed model of mental illness endophenotypes (5,40). In this model, stress reactivity, lack of self-control, and reward sensitivity were assumed to each contribute to the Uncontrolled Eating phenotype.

Finally, we examined the relationships between Uncontrolled Eating, the impulsivity traits, BMI and longitudinal changes in BMI in a hierarchical four-dimension model (Model 4, Figure 1D). We tested whether BMI and longitudinal change in BMI (delta BMI) were predicted by Uncontrolled Eating, stress reactivity, lack of self-control and reward sensitivity.

We used the Lavaan package in R version 3.3. (41) to perform SEM. Model fit was assessed with the chi-square test (X^2^), the Comparative Fit index (CFI), Root Mean Square Error of Approximation (RMSEA) and the standardized root mean squared residuals (SRMR). The following guidelines were utilized for judging good fit: RMSEA (acceptable fit < 0.08, good fit < 0.05), CFI (acceptable fit 0.95-0.97, good fit > 0.97), SRMR (acceptable fit 0.05-0.10, good fit < 0.05) (42).

To test for the robustness of the preferred hierarchical SEM model (model 3, Figure 1C), we repeated the analysis by randomly splitting the sample in half. Age and sex were regressed out of all the observable variables in the models. To account for the multiple models performed, we set a stringent p value threshold (p<0.005).

### Magnetic Resonance Imaging parameters and preprocessing

High-resolution T1-weighted anatomical images with voxel size = 1×1×1 mm were obtained (TR = 2.3 s; TE = 2.98 ms; FOV phase=93.8°; FOV = 256mm) with a Siemens Magnetom Trio 3T MRI scanner at the Montreal Neurological Institute (MNI).

Pre-processing of T1-weighted MRIs included denoising using optimized non-local means filtering (43), correction for intensity inhomogeneity (44) and linear intensity scaling using histogram matching to the ICBM-MNI152 template. The images were linearly registered to the ICBM-MNI152 template (9 parameter registration) (45). A mask of the brain was generated using BEaST, a nonlocal segmentation method applied to the linearly registered images in stereotaxic space (46).

### Cortical Thickness

All T1-weighted MRI images were processed using the CIVET pipeline (version 2.1; http://www.bic.mni.mcgill.ca/ServicesSoftware/CIVET). Native T1-weighted MRI scans were first corrected using the N3 algorithm, underwent brain masking, and registration to ICBM-MNI152 template. Images were then segmented into gray and white matter, cerebrospinal fluid and background with a neural net classifier. The white matter (inner) and gray matter (outer) cortical surfaces were generated using the Constrained Laplacian-based Automated Segmentation with Proximities algorithm. These surfaces were resampled to a stereotaxic surface template to allow vertex-based measurement of cortical thickness. All resulting images went through a stringent quality control by two inspectors in which 69 of 71 images were accepted for further analysis. Cortical thickness in MNI space was defined as the linked distance between the two surfaces across vertices.

### Post-hoc brain-impulsivity analysis

We performed a post-hoc exploratory analysis to test which brain regions are associated with the three general-impulsivity domains. We extracted the latent individual scores for stress reactivity, lack of self-control and reward sensitivity of the participants with an available MRI. Next, using the SurfStat software package (http://www.math.mcgill.ca/keith/surfstat/), we computed three separate linear regression models to test for the associations between the three latent impulsivity scores and cortical thickness. The models were analyzed with random field theory with a threshold of p<0.05 corrected for multiple comparisons over the entire surface (47). Results are reported cluster- and vertex-wise corrected. Total surface area was included as covariate. Age and sex were already regressed out of the observable variables forming the latent scores (see Structural Equation Models above). For this reason, these confounding variables were not included in the analyses. Moreover, since the latent scores were obtained from a SEM model that includes the 3 general impulsivity variables as latent variables, each latent score “accounts” for the other two.

## Results

Participants’ characteristics are listed in Table 1.

### Uncontrolled Eating and general impulsivity traits can be considered independent latent variables (Models 1 and 2)

A four-dimension model, with Uncontrolled Eating, stress reactivity, lack of self-control and reward sensitivity as latent variables provided an acceptable fit (χ^2^_(29)_ = 322.40; RMSEA = 0.068; CFI= 0.956; SRMR = 0.037). The first latent factor, Uncontrolled Eating, had positive loadings from Disinhibited Eating, Power of Food, and Emotional Eating. The latent factor stress reactivity had a negative loading from the Rosenberg self-esteem questionnaire and positive loadings from Neuroticism and the Perceived Stress Scale. Lack of self-control, the third latent factor had a positive loading from Lack of Perseverance and a negative loading from Conscientiousness. Reward sensitivity was characterized by positive loadings from the Extraversion and Sensation Seeking (UPPS) subscales. This model provided a better fit than an alternative single-dimension model, where all the observable variables were fit into a single latent score (difference in χ^2^_(6)_ = 2770.5; p=2.2e-16).

### A four-dimensional hierarchical model, where general impulsivity traits predict Uncontrolled Eating, is plausible (Model 3)

We tested the plausibility of a four-dimension hierarchical model. In this model, the latent variables stress reactivity, lack of self-control and reward sensitivity were characterized as predictors of Uncontrolled Eating. All the paths were significant. Note that the fit parameters are the same as Model 2, since sample size and degrees of freedom did not change and provided an acceptable fit (Figure 2A).

**Figure 2.**
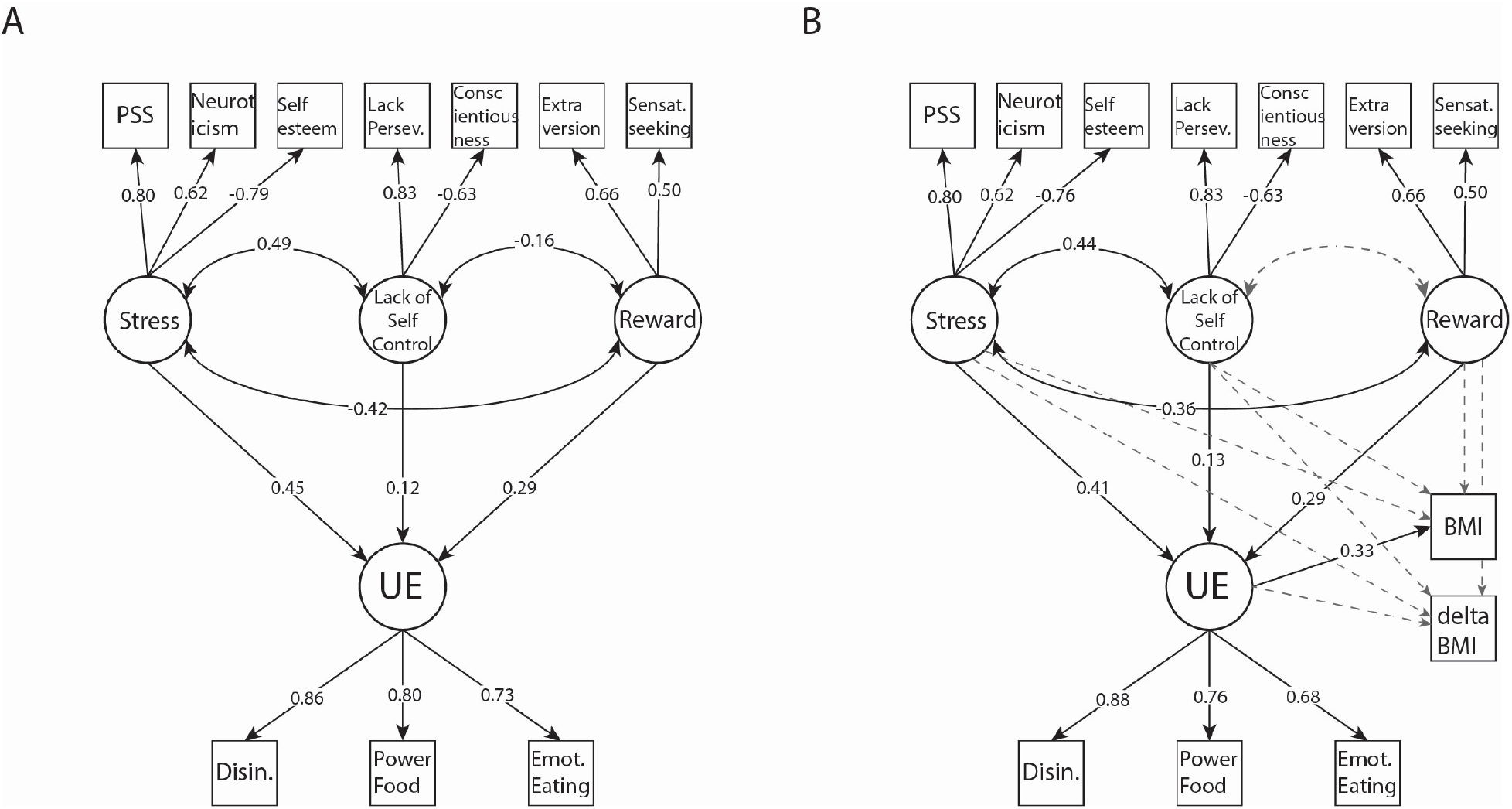
SEM representing: A) Model 3 (n=2318), the relationships between Uncontrolled Eating (UE) and the general impulsivity-related variables stress reactivity, lack of self-control and reward sensitivity. B) Model 4 (n=1197), a model that additionally includes the associations between UE, general impulsivity-related variables, BMI and 6-month longitudinal change in BMI (delta BMI) are added. Numeric values are standardized beta weights (all of them p<0.005). Dashed gray lines depict non-significant associations. PSS – perceived stress scale. UE – Uncontrolled Eating. BMI – body mass index. Stress – stress reactivity. Reward – reward sensitivity. Disinh. – disinhibition. Emot. Eating – emotional eating. Lack Persev. – lack of perseverance. Sensat. seeking – sensation seeking.

### Relationship between Uncontrolled Eating, BMI and longitudinal weight change (Model 4)

Finally, we tested the relationship between Uncontrolled Eating, general impulsivity, BMI and longitudinal changes in BMI in the hierarchical four-dimension model. From these additional regression paths, the only one that was significant was the association between BMI and Uncontrolled Eating (B=0.38; SE=0.05; Beta=0.09; p<0.001). This model provided an acceptable fit (χ^2^_(41)_ = 198.52; RMSEA = 0.055; CFI= 0.956; SRMR = 0.032) (Figure 2B).

### Validation of Model 4 in the half split dataset

All the previously observed relationships in Model 4 remained equivalent in the two subsamples. Fit parameters in the first half of the sample (N=601 observation) were as follows: (χ^2^_(41)_ = 102.97; RMSEA = 0.050; CFI= 0.965; SRMR = 0.032). The same model in the second half of the sample (N=600 observations) had the following fit parameters (χ^2^_(41)_ = 142.72; RMSEA = 0.064; CFI= 0.939; SRMR = 0.039).

### Post-hoc correlations between cortical thickness and general impulsivity scores

Higher stress sensitivity was associated with greater cortical thickness in the right temporoparietal junction extending to superior temporal gyrus. Lack of self-control was positively associated with greater thickness in the right superior frontal gyrus extending to midcingulate cortex. Finally, higher scores in reward sensitivity were associated with lower thickness in the bilateral inferior frontal gyrus (Figure 3).

**Figure 3.**
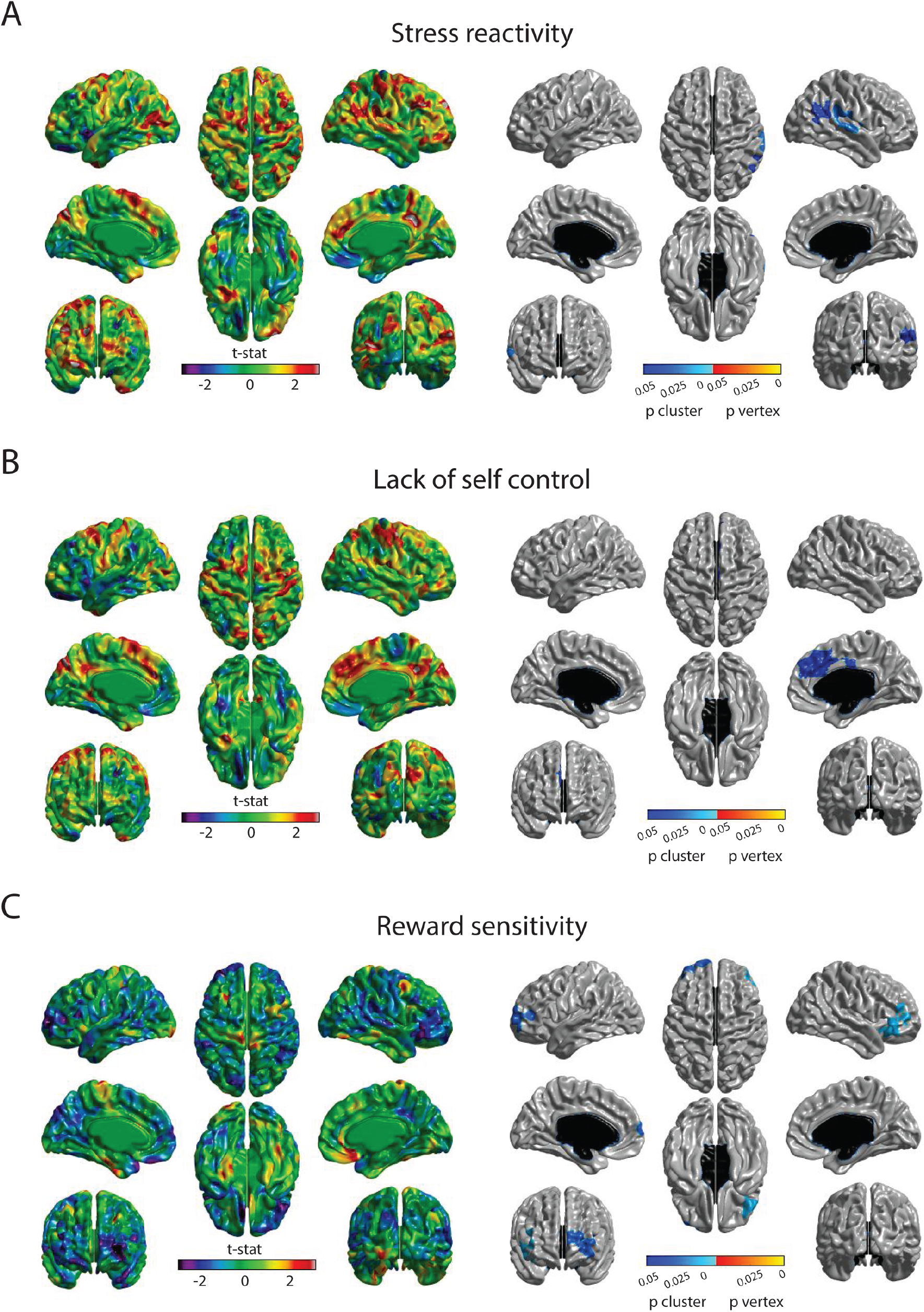
Correlations between latent factors of impulsivity and cortical thickness.

## Discussion

The purpose of the current study was to provide a comprehensive examination of the relationship between domain-general (personality-related facets of impulsivity) and eating-specific obesity endophenotypes (Uncontrolled Eating), BMI, and weight changes. To do so, we tested a series of structural equation models in a large sample size (N>2300) of first-year students and accounted for the multidimensional nature of impulsivity. A final SEM model, which depicted a hierarchical relationship between general impulsivity, Uncontrolled Eating, BMI and weight gain, was developed in the first split-half of the sample and replicated in the second. In a follow-up analysis we also investigated neural correlates of different facets of impulsivity. We found that cortical thickness in parts of the frontal, cingulate and temporoparietal cortex were correlated with impulsivity and hence are potentially related to BMI via Uncontrolled Eating.

Following previous work, we first stratified impulsivity into three general latent variables: stress sensitivity, lack of self-control and reward sensitivity (8,10). A fourth latent variable, Uncontrolled Eating, was considered an eating-specific form of impulsivity. In line with previous publications (27), Uncontrolled Eating was defined using the total scores of relevant eating questionnaires. We showed that an SEM that separated these 4 latent factors was significantly better than a model in which all the observable variables were loaded into a single latent score. Moreover, a hierarchical relationship, in which the three general impulsivity domains were considered predictors of Uncontrolled Eating, was found to be plausible. Each general impulsivity domain had an independent and positive correlation with Uncontrolled Eating. We also found that higher scores in Uncontrolled Eating were positively associated with BMI, as demonstrated previously (27,28). However, none of the relations between general impulsivity domains and BMI were significant in the model that also included Uncontrolled Eating. The SEM model therefore suggests that the association between impulsivity and BMI is indirect, with impulsivity factors predicting Uncontrolled Eating, and Uncontrolled Eating predicting BMI.

Finally, we tested the hypothesis that Uncontrolled Eating underlies vulnerability for weight gain during the first year of university. We utilized the transition period into university because it is associated with a high risk of weight gain. In our sample, the average weight gain was small (0.52 kg over 10 months), but significant, and was similar to previous reports (29). However, in this sample we did not find evidence that Uncontrolled Eating can longitudinally predict weight gain. This could be due to limited duration of the study. Moreover, we cannot exclude the possibility of seasonal weight gain (48), which would not necessarily be related to Uncontrolled Eating. The existence of bidirectional relationships between Uncontrolled Eating and BMI might also mask a relationship, since changes in body weight might lead to longitudinal variations in eating-specific impulsivity (49). For instance, scores in different Uncontrolled Eating scales decrease after bariatric surgery (50) or voluntary weight loss (51).

Our results support a model in which BMI is associated with Uncontrolled Eating, which in turn is derived from general impulsivity. Impulsivity could compromise the maintenance of behaviors that promote healthy weight. High scores on conscientiousness have been consistently linked to a lower risk of having and developing obesity across independent samples (52,53). These results are generally derived from sample populations with a higher mean age than ours, suggesting that self-control deficits may be more associated with BMI as individuals age and start making their own food decision. Nonetheless, our findings point to reduced conscientiousness (lack of self-control) as a risk factor for overeating behaviors in adolescent and young adult populations.

In a subset of the sample, we performed additional post-hoc correlations between cortical thickness and impulsivity in all three tested domains. We found that the temporoparietal junction, superior frontal gyrus/midcingulate cortex and inferior frontal gyrus were associated with stress sensitivity, lack of self-control and reward sensitivity, respectively. These results are consistent with previous reports showing brain correlates of impulsivity (19,20,54) and partially overlap with obesity-related brain regions (21,55). Paralleled by our SEM results, these findings show that cortical thickness differences might indirectly influence BMI via Uncontrolled Eating. In the future we would like to update our SEM model to test the hypothesis that brain differences can act as independent predictors of impulsivity, which in turn are significant predictors of Uncontrolled Eating, and hence BMI. This hierarchy of relationships can be depicted as a watershed model (40), which will allow to test the hypothesis that the temporoparietal junction, midcingulate cortex and inferior frontal gyrus are indirectly involved in BMI, via their association with impulsivity. Unfortunately, due to low imaging sample size, this was not possible in this study.

The results of our study should be considered with regards to its limitations. Although we had a large sample group, whereby we were able to detect and replicate results from meta-analyses, most of our measures were self-reported. In addition, brain imaging was conducted with a small subset of the total sample.

## Acknowledgments

This research was supported by a Canadian Institutes of Health Research Grant to AD. IGG was the recipient of a Postdoctoral Fellowship from the Canadian Institutes of Health Research. SN was supported by a Frederick Banting and Charles Best Canada Graduate Scholarship. UV was supported by Personal Post-doctoral Research Funding project PUTJD654 and by Fonds de recherche du Québec – Santé (FRQS) foreign post-doctoral training award.

## Competing interests

IGG, SN, FM, MD, YY, SGS, YZ, NS, DLC, UV, AD report no financial interests or potential conflicts of interest.

## Notes

### Competing Interest Statement

The authors have declared no competing interest.

